# The structure of the genetic code as an optimal graph clustering problem

**DOI:** 10.1101/332478

**Authors:** Paweł Błażej, Dariusz R. Kowalski, Dorota Mackiewicz, Małgorzata Wnetrzak, Daniyah A. Aloqalaa, Paweł Mackiewicz

## Abstract

The standard genetic code (SGC) is the set of rules by which genetic information is translated into proteins, from codons, i.e. triplets of nucleotides, to amino acids. The questions about the origin and the main factor responsible for the present structure of the code are still under a hot debate. Various methodologies have been used to study the features of the code and assess the level of its potential optimality. Here, we introduced a new general approach to evaluate the quality of the genetic code structure. This methodology comes from graph theory and allows us to describe new properties of the genetic code in terms of conductance. This parameter measures the robustness of codon groups against the potential changes in translation of the protein-coding sequences generated by single nucleotide substitutions. We described the genetic code as a partition of an undirected and unweighted graph, which makes the model general and universal. Using this approach, we showed that the structure of the genetic code is a solution to the graph clustering problem. We presented and discussed the structure of the codes that are optimal according to the conductance. Despite the fact that the standard genetic code is far from being optimal according to the conductance, its structure is characterised by many codon groups reaching the minimum conductance for their size. The SGC represents most likely a local minimum in terms of errors occurring in protein-coding sequences and their translation.

## Introduction

The standard genetic code (SGC) is simply the set of rules according to which the information stored in DNA molecule can be transmitted into the protein world. This code is nearly universal for three domains of life, Bacteria, Archaea and Eukaryota, which means that almost all living organisms decode their genes into proteins on the same basis. The code uses 64 nucleotide triplets, called codons, to encode 20 amino acids and stop translation signal. Since the number of amino acids is smaller than the number of codons and each codon has to code any information, the SGC must be degenerated, i.e. there exists an amino acid that is encoded by more than one codon, i.e. a group of codons. The redundant codons, called synonymous, are organized in specific groups. Nine amino acids are encoded by groups of two codons, called twofold degenerated. Five amino acids have codons that are fourfold degenerated, and three amino acids have six codons. One amino acid is coded by three codons, and only two amino acids, i.e. methionine and tryptophan, have single codons. Three codons, called stop codons, break the synthesis of proteins in the translation process.

This degeneracy of the genetic code has puzzled biologists since the code was cracked [Khorana et al., 1966, Nirenberg et al., 1966]. One explanation of this phenomenon was suggested by Francis Crick, who assumed that only the first two codon positions were important in a primordial code [Crick, 1968]. Some evidence for this hypothesis is in the way of decoding information by transfer RNA (tRNA) during the protein translation process. Each tRNA decodes a codon by a complementary triplet, called an anticodon, and carries a single amino acid that matches this codon in the transcript (mRNA). However, it is not necessary for each codon to have its corresponding anticodon because one tRNA can decode more than one codon. The ambiguity of this recognition results from the less specific interactions between base pairs in the first anticodon position and the third codon position, which is explained by the Wobble Hypothesis [Crick, 1966]. The lesser specificity is often associated with the post-transcriptional modifications of the nucleotide at the first position of the anticodon in tRNA [Murphy and Ramakrishnan, 2004]. In consequence, the base in the first anticodon position can pair with more than one base type in the third codon position. For example, a nucleoside inosine, derived from the modified adenine, can recognize even three bases, adenine, cytosine and uracil. Moreover, some aminoacyl-tRNA synthetases, i.e. specific enzymes, which charge an amino acid to the appropriate tRNA, recognize only the last two nucleotide bases of the anticodon to decide which amino acid to attach [Fukai et al., 2003, Sankaranarayanan et al., 1999, Yaremchuk et al., 2000]. Thus, the first two bases of the codon play a more important role in the specific codon-anticodon recognition than the third codon position.

There is an interesting consequence of the genetic code redundancy related to the mutation process. The substitution of one nucleotide to another in the degenerated codon positions does not change the coded amino acid. Such types of mutations are called synonymous or silent, whereas those that change the coded information, amino acid or stop signal, are named nonsynonymous. The degeneracy implies a specific structure and properties of the genetic code in terms of these mutations. It is evident that this property can also have a decisive impact on the potential robustness of the genetic code against amino acid and stop signal replacements. The proper structure of the code associated with the degeneracy can minimize the number of these replacements. Such properties were noticed in the standard genetic code and it was suggested that the code could have evolved to minimize the consequences of translational errors and substitutions in protein coding sequences [Ardell, 1998, Ardell and Sella, 2001, B lażej et al., 2016, Di Giulio, 1989, Di Giulio and Medugno, 1999, Epstein, 1966, Freeland and Hurst, 1998b, Freeland and Hurst, 1998a, Freeland et al., 2003, Freeland et al., 2000, Gilis et al., 2001, Goldberg and Wittes, 1966, Goodarzi et al., 2005, Haig and Hurst, 1991, Woese, 1965].

Since the genetic code is a set of codons which are related, e.g. by nucleotide substitutions, the general structure of this code can be well described by the methodology taken from graph theory [Beineke and Wilson, 2005, Lee et al., 2014]. Similarly to [Tlusty, 2010], we assume that the code encodes 21 items, i.e. 20 amino acids and stop translation signal, and all 64 codons create the set of vertices of a graph, in which the set of edges corresponds to all possible single-nucleotide substitutions occurring between the codons. In this representation, each genetic code is a partition of the set of vertices into 21 disjoint subsets. Therefore, the optimization problem of the genetic code in regard to the substitutions can be reformulated into the optimal graph clustering problem. From the computational point of view, this problem is NP-hard. Obviously, there are various approximation methods but in the general case they need also a substantial computational effort and do not guarantee finding the optimal solution.

To study the optimality of the general structure of the genetic code, we modified the set conductance measure, which is widely used in graph theory [Lee et al., 2014] and has many practical interpretations, for example in the theory of random walks [Levin et al., 2009] and social networks [Bollobás, 1998]. In the problem considered here, the conductance of a codon group is the ratio of nonsynonymous substitutions to all possible single nucleotide substitutions in which the codons in this group are involved. Therefore, this parameter can be interpreted as a measure of robustness against the potential changes in protein-coding sequences generated by the single nucleotide substitutions. Moreover, we also defined the minimum *k*-set conductance evaluated from all sets of vertices with a fixed size *k*. Based on these definitions, we introduced two different characteristics of genetic codes quality. The first one, called the code maximum conductance, describes a given genetic code in terms of the maximum set conductance value calculated for its codon groups. The second one is the average conductance value calculated as an arithmetic mean of codon group conductance. Using this methodology, we found some exact solutions, i.e. the optimal genetic codes, in respect to the postulated measures.

## Preliminaries

To study the general structure of the genetic code we developed its graph representation. Let *G*(*V, E*) be a graph in which *V* is the set of vertices representing all possible 64 codons, whereas *E* is the set of edges connecting these vertices. All connections fulfill the property that the vertices, i.e. codons *u, v ∈ V* are connected by the edge *e*(*u, v*) *∈ E* (*u ∼ v*) if and only if the codon *u* differs from the codon *v* in exactly one position. From the biological point of view, the edges represent all possible single nucleotide substitutions, which occur between codons in a DNA sequence. In our case, we claim that all nucleotide substitutions are equally probable to avoid arbitrary assumptions on the mutational process. Hence, the graph *G* is undirected, unweighted and regular with the vertices degree equal to 9.

Following graph theory, each potential genetic code *𝒞* which codes 20 amino acids and stop translation signal is a partition of the set *V* into 21 disjoint subsets, i.e. groups of codons, *S*. Thus, we obtain the following representation of genetic code *𝒞*:

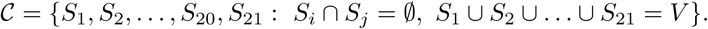

In Fig. 1 we showed an example of the partition of the graph *G* which corresponds to the standard genetic code. Many properties of the genetic code strongly depend on the types and the number of connections between the groups of codons. From the biological point of view, it is interesting to study the code structure according to the number of connections between and within the codon groups. These connections correspond to nonsynonymous and synonymous substitutions, respectively. The code that minimizes the number of the nonsynonymous substitutions is regarded the best because it decreases the biological consequences of mutations [Ardell, 1998, Di Giulio, 1989, Freeland and Hurst, 1998b, Freeland and Hurst, 1998a, Freeland et al., 2003, Haig and Hurst, 1991, Woese, 1965]. Therefore, the conditions under which the partitions of the graph vertices describe the best genetic code, are worth finding.

**Figure 1.**
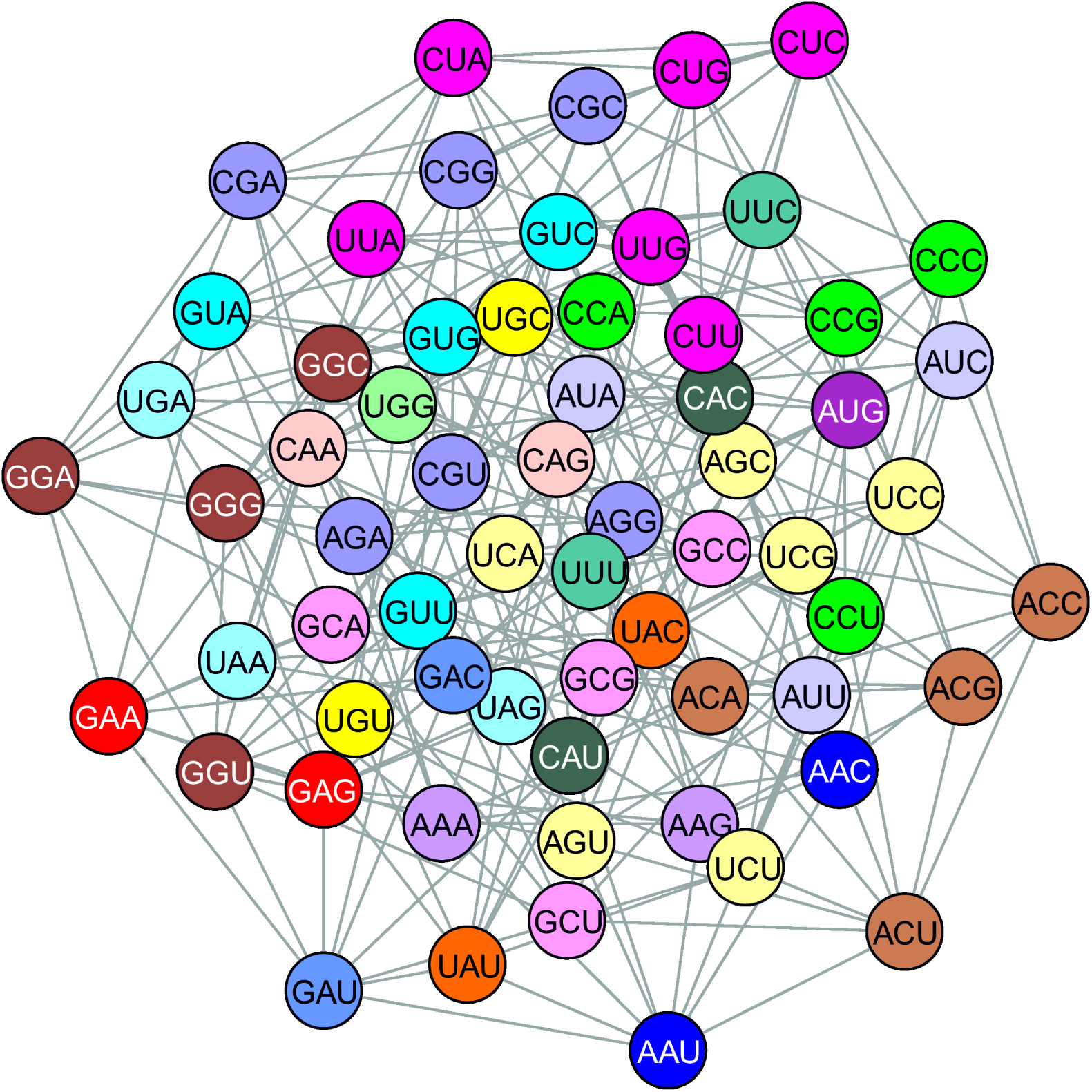
The standard genetic code as an example of the partition of the graph *G*(*V, E*). Every group of vertices with the same colour corresponds to the respective set of codons which code the same amino acid or stop translation signal. The edges represent all possible single nucleotide substitutions.

There are many methods of the optimal graph partitioning, which are based on different approaches. In this work, to investigate the theoretical features of genetic codes in terms of connections between the codon groups, we decided to use the set conductance measure, which plays a central role in the spectral graph clustering method. The definition of the set conductance measure is as follows:

#### Definition 1.

For a given graph G let S be a subset of V. The conductance of S is defined as:

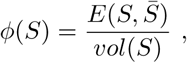

where 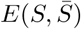 is the number of edges of G crossing from S to its complement 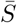 and vol(*S*) is the sum of all degrees of the vertices belonging to S.

The set conductance has several interpretations. For example, in the theory of random walks *ϕ*(*S*) is the probability that a simple random walk, which starts at a random vertex of *S*, leaves this set in one step. This observation is a good starting point to define the optimality of a given codon group which encodes an amino acid.

The definition of the set conductance allows us to define the maximum conductance of a genetic code:

#### Definition 2.

The maximum conductance of a genetic code 𝒞 is defined as:

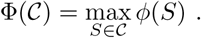

The measure Φ(*𝒞*) provides an important information about the general properties of the genetic code and the codon groups because it characterizes the worst codon group in terms of set conductance. The definition of the best code Φ_*min*_ results in a natural way and is given by the formula:

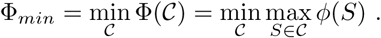

The definition of Φ_*min*_ is similar to the definition of the *k*-th order graph conductance [Lee et al., 2014] and has a useful interpretation because if we assume that the value of Φ_*min*_ is small then there exists the partition of the graph *G*, i.e. the genetic code, in which each codon group has a small set conductance. Therefore, it gives us the lower bound of the genetic code robustness against changes in the translation of protein-coding sequences.

Besides the maximum conductance it is also interesting to calculate the average conductance of a given code. This measure gives us a more general view of the genetic code properties and is realized by the following definition:

#### Definition 3.

The average conductance of a genetic code C is defined as:

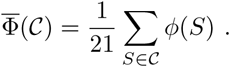

Using the definition presented above, we are able to describe the best code in terms of the average conductance, which is defined as follows:

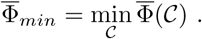

Similarly to the definition of Φ_*min*_, 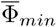 gives us a lower bound of the genetic code robustness measured in terms of the average code conductance.

It seems reasonable to claim that the optimal codon group should have a low set conductance which means that the number of nucleotide substitutions that change the translation of the protein-coding sequence is relatively low in comparison to the total number of all possible nucleotide changes. In this context, it is also interesting to calculate the *k*-size-conductance *ϕ*_*k*_(*G*) described as the minimal set conductance over all subsets of *V* with the fixed size *k*.

### Definition 4.

The k-size-conductance of the graph G, for k ≥ 1, is defined as:

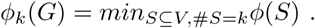

Calculating *ϕ*_*k*_(*G*) for the fixed *k* and establishing its correspondence to a codon group is crucial from the biological point of view because the codon group with the minimal *k*-size-conductance seems to be the most robust against changes in the translation of protein-coding sequences. What is more, the definition of the *k*-size-conductance is a good starting point for further investigation of the whole space of all possible genetic codes. To do so, we introduce two subsequent definitions.

### Definition 5.

Let κ be a vector of integers that fulfills the following properties:

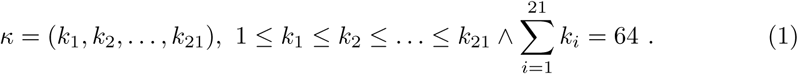

Using the Definition 5, we get an immediate conclusion that for every genetic code *𝒞*, there exists a vector of integers *κ*_*C*_ which satisfies (1) and represents a sequence of codon group sizes in the ascending order. What is more, it is possible to split the whole set of all possible genetic codes into equivalence classes [*κ*] where:

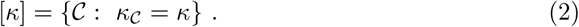

Using this characterization, we can formulate the definition of the average *κ*-size-conductance.

### Definition 6.

Let κ be a vector of integers such that the condition (1) holds and let [κ] be an equivalence class defined by (2), then the average κ-size-conductance Φ[κ] is described as:

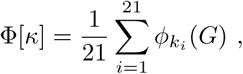

where κ = (k_1_, k_2_, …, k_21_).

It is evident that using the Definition 6 we get a lower bound of the average code conductance for every genetic code *𝒞*. This fact is pointed up in the next proposition.

### Proposition 1.

Let 𝒞 be a genetic code such that𝒞 ∈ [κ], then the following inequality holds:

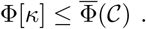

*Proof*. It is an immediate conclusion from the definition of the k-size-conductance.

It should be noted that the problem of finding the set of edges *S, S* ⊆ *V* which satisfies #*S* = *k* and *ϕ*(*S*) = *ϕ*_*k*_(*G*) is NP-complete for graphs in general. Fortunately, the graph *G*, describing interactions between codons generated by single nucleotide substitutions, has some desirable properties, which allow us to generate the sets of vertices *S* with the fixed size #*S* = *k* and *ϕ*(*S*) = *ϕ*_*k*_(*G*) quite easily. This method of fast establishing the optimal sets in respect to *ϕ*_*k*_(*G*) results from two subsequent propositions:

### Proposition 2.

Graph G can be represented as a Cartesian graph product K_4_ × K_4_ × K_4_, where K_4_ is 4-clique with the set of vertices {A, U, G, C}. Moreover, two vertices (x, y, z), (x^′^, y^′^, z^′^) are connected by the edge e((x, y, z), (x^′^, y^′^, z^′^)) if (x = x^′^ and y = y^′^ and z ∼ z^′^) or (x = x^′^ and y ∼ y^′^ and z = z^′^) or (x ∼ x^′^ and y = y^′^ and z = z^′^).

The next proposition gives us a very useful characterization of a set of vertices reaching *k*-size-conductance from all possible subsets with a given size *k*.

### Proposition 3.

Let us consider a linear order of the set of vertices of 4-clique K_4_, A > C > G > U, and let F(k) be the collection of the first k vertices of a graph K_4_ × K_4_ × K_4_ = G in the lexicographic order, then we get:

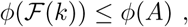

where A ⊆ K_4_ × K_4_ × K_4_, #A = k, for any k ≥ 1. Therefore, the following equations hold for any k ≥ 1:

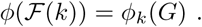

This proposition is an immediate conclusion from the Theorem 1 presented in the paper [Bezrukov and Elsässer, 2003], where the authors dealt with the edge-isoperimetric problem of the Cartesian powers of graphs. This question can be reformulated to the problem of finding *ϕ*_*k*_(*G*) for *k* ≥ 1. As a consequence, we get a nice and efficient method for calculating *ϕ*_*k*_(*G*), which is described in the following proposition:

### Proposition 4.

Let A = (a_ij_) be an adjacency matrix of a graph G, where rows and columns are sorted in the lexicographic order, then the first k ≥ 1 vertices create a set with the set conductance equal to the k-size-conductance ϕ_k_(G). Then, ϕ_k_(G) can be calculated according to the formula:

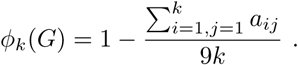

In the Table 1, we show the example of the upper-left *k* by *k* submatrix extracted from the adjacency matrix of graph *G* (shown in Electronic Supplementary Material ESM 1). Applying Proposition 4 to this example, we are able to calculate the *k*-size-conductance *ϕ*_*k*_(*G*) of subgraphs for *k* = 1, 2, …, 9 (Fig. 2), which will be useful later in the analysis of the minimum average conductance of genetic codes.

**Table 1.** The example of upper-left *k* by *k* submatrix extracted from the graph *G* adjacency matrix of codons, where rows and columns are ordered in the lexicographical order. In the light of the Proposition 3, the presented submatrix allowed us to calculate *ϕ*_*k*_(*G*) and to determine the structures of *ϕ*_*k*_(*G*), i.e. the optimal subgraphs for *k* = 1, 2, … 9. The full matrix is presented in Electronic Supplementary Material ESM_1.

|  | AAA | AAC | AAG | AAU | ACA | ACC | ACG | ACU | AGA |
| --- | --- | --- | --- | --- | --- | --- | --- | --- | --- |
| AAA | 0 | 1 | 1 | 1 | 1 | 0 | 0 | 0 | 1 |
| AAC | 1 | 0 | 1 | 1 | 0 | 1 | 0 | 0 | 0 |
| AAG | 1 | 1 | 0 | 1 | 0 | 0 | 1 | 0 | 0 |
| AAU | 1 | 1 | 1 | 0 | 0 | 0 | 0 | 1 | 0 |
| ACA | 1 | 0 | 0 | 0 | 0 | 1 | 1 | 1 | 1 |
| ACC | 0 | 1 | 0 | 0 | 1 | 0 | 1 | 1 | 0 |
| ACG | 0 | 0 | 1 | 0 | 1 | 1 | 0 | 1 | 0 |
| ACU | 0 | 0 | 0 | 1 | 1 | 1 | 1 | 0 | 0 |
| AGA | 1 | 0 | 0 | 0 | 1 | 0 | 0 | 0 | 0 |

**Figure 2.**
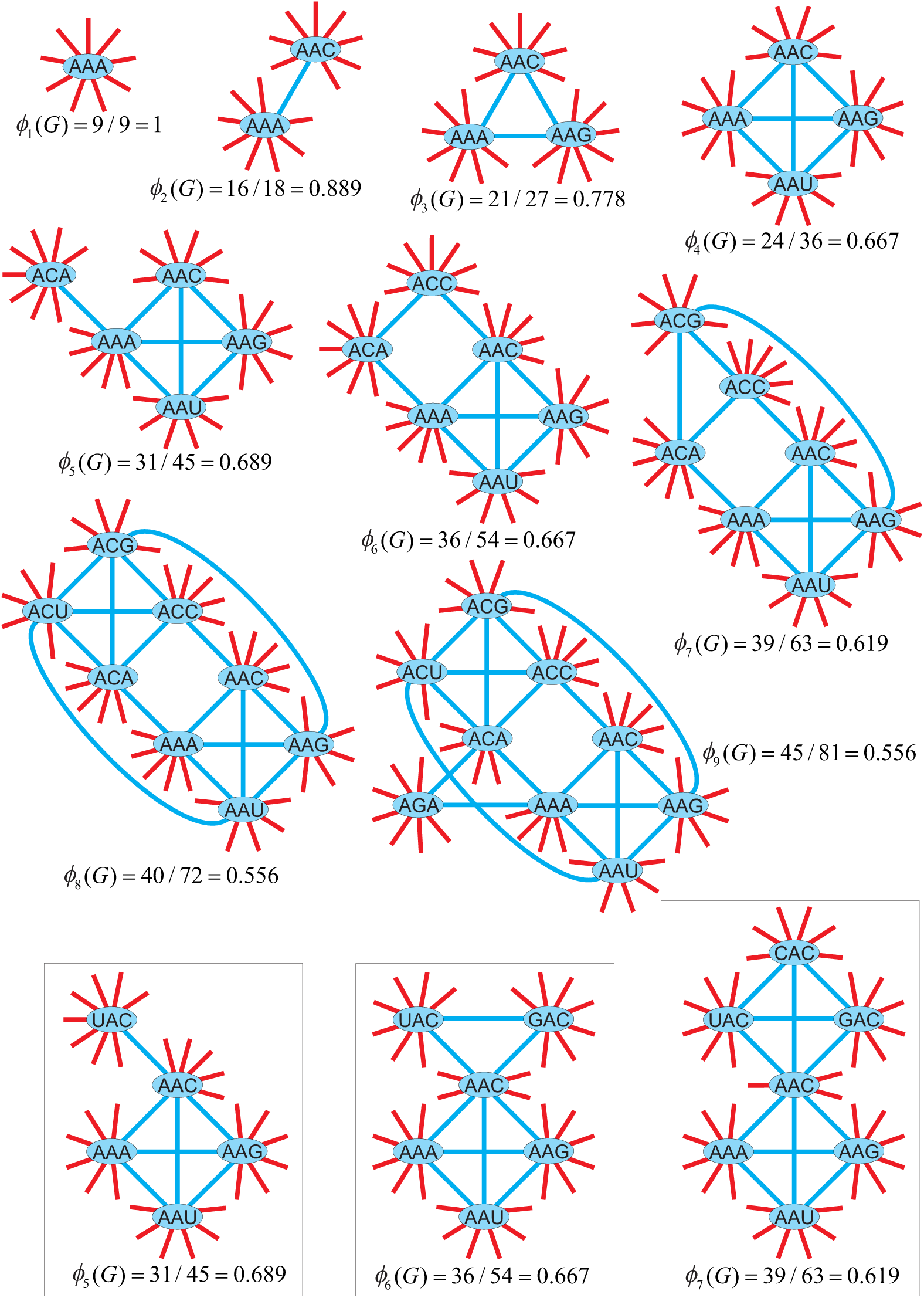
The examples of the codon subgraphs optimal in terms of the *k*-size-conductance, with the number of vertices *k* from 1 to 9. The calculation of its *k*-size-conductance *ϕ*_*k*_(*G*) is shown below the given subgraph. The red lines mean nonsynonymous substitutions and the blue ones indicate synonymous substitutions. Three subgraphs outlined with boxes represent alternatives for *k* = 5, 6 and 7.

## 1 Results and Discussion

### 1.1 The conductance of codon groups with different size

The main goal of our work is to find the optimal genetic codes in terms of two characteristics, the maximum conductance Φ_*min*_ and the average conductance 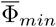. Furthermore, we compare the properties of these codes with the standard genetic code, which is interesting from the biological point of view. Using the Proposition 4, we calculated the *k*-size-conductance *ϕ*_*k*_(*G*) for *k* = 1, 2, …, 64, i.e. groups consisting of different number of codons. The *ϕ*_*k*_(*G*) values calculated for 1 ≤ *k* ≤ 9 are presented in Fig. 2 with corresponding subgraphs. It is interesting that *ϕ*_*k*_(*G*) reaches the same values for *k* = 4, 6 and *k* = 8, 9.

The relationship between the *k*-size-conductance *ϕ*_*k*_(*G*) and the *k* size of all codon groups is plotted in Fig. 3. As expected, the values of *ϕ*_*k*_(*G*) decrease with the growth of the set size *k*. Particularly, *ϕ*_*k*_(*G*) declines rapidly from *k* = 1 to *k* = 16 then starts to decrease gradually till *k* = 64. Interestingly, there are some local minima for *k* = 4, 8, 16, 32 and 48 in the general decreasing trend. This fact suggests some interesting connections between the structures of *ϕ*_*k*_(*G*)-optimal subgraphs of the graph *G*.

**Figure 3.**
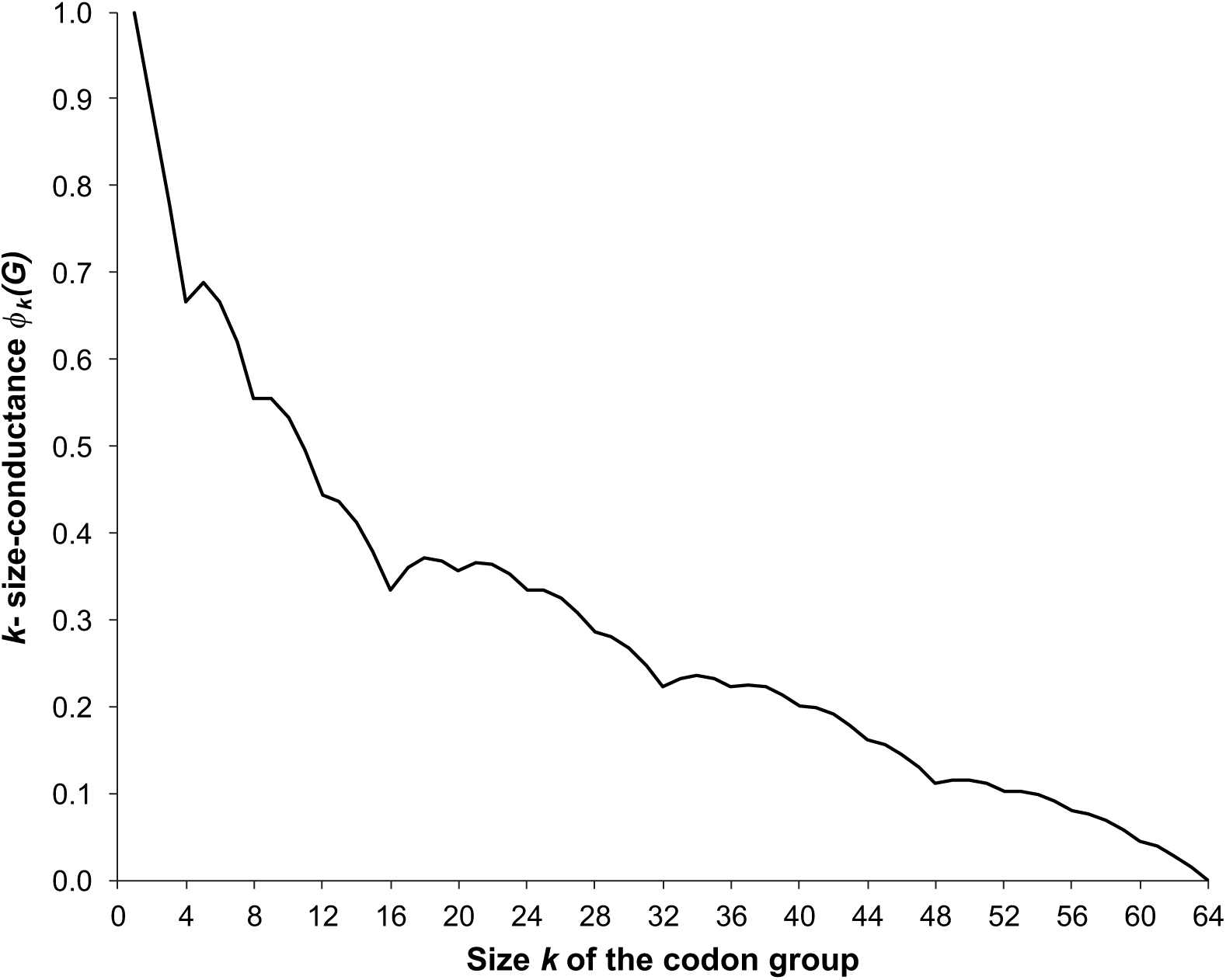
The relationship between the *k*-size-conductance *ϕ*_*k*_(*G*) and the *k* size of codon groups.

### 1.2 The optimal genetic code in respect to the maximum code conductance

As was stated in the Preliminary section, the task of finding the optimal genetic code can be reformulated as the question about the optimal graph partition. However, this is a well-known NP-hard problem for graphs in general, which implies that obtaining the potential solution is time consuming and requires substantial computational effort. To avoid this inconvenience, many approximate methods have been developed but generally they are able to find only near-optimal results. Fortunately, we found the exact solution, i.e. the optimal genetic code in respect to the minimum of maximum conductance of the genetic code, without complicated calculations or advanced theoretical methodology. Our investigation was based on several simple observations related to the properties of the graph *G* and the general features of the genetic code.

To describe the optimal code in terms of the minimization of the code conductance, it is enough to consider the following facts:

#### Lemma 1.

The maximum conductance of a code 𝒞 is not smaller than the k-size-conductance of the subset with the minimal size k, that is:

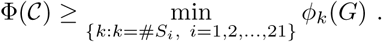

*Proof.* Let us consider a graph partition *S*_1_, *S*_2_, *…, S*_21_ which corresponds to *𝒞* and let *S*_*k*_ be a codon group with the smallest size. Hence, we get immediately that Φ(*𝒞*) *≥ ϕ*(*S*_*k*_) and also *ϕ*(*S*_*k*_) *≥ ϕ*_*k*_(*G*) by the definition of the *k*-size-conductance *ϕ*_*k*_(*G*), which proves the proposition.

The next lemma is related to the size of codon groups and the number of items, i.e. amino acids and stop translation signal.

#### Lemma 2.

If the genetic code 𝒞 encodes 20 amino acids and stop translation signal, then there exists a set in its graph partition that contains fewer than four codons.

*Proof.* It is an obvious remark, because otherwise the code 𝒞 would code at most 16 amino acids.

Using Lemmas 1 and 2, we are able to give the lower bound of the maximum conductance value of the best genetic code.

#### Lemma 3.

The maximum conductance of the optimal genetic code fulfills the following formula:

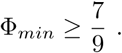

*Proof.* This proof is the immediate consequence of Lemma 1 and 2. Since the optimal code has at least one codon group consisting of fewer than 4 codons, then depending on the minimal size of this group, the code conductance is not smaller than *ϕ*_1_(*G*), *ϕ*_2_(*G*) or *ϕ*_3_(*G*). Out of these values, the minimal one is 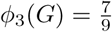, which gives us the lower bound of Φ_*min*_.

Studying the genetic codes in which an amino acid is coded by more than 4 codons leads to the following observation.

#### Lemma 4.

If the genetic code 𝒞 has a codon group with the size greater than 4, then its maximum conductance fulfills the following inequality:

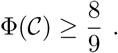

*Proof.* Let us assume that there exists a codon group consisting of five codons in the given code. Thereby, we have to create the 20-sets partition using 59 codons. Thus, it is impossible to create 20 subsets, each consisting of three codons. Therefore, using Lemma 1 and the inequality:

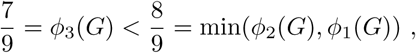

we complete the proof of this lemma.

Moreover, using the method presented in the proof of Lemma 4, we can easily show that the optimal code cannot include more than one codon group with the size *k* = 4.

To sum up all the facts presented above, we can formulate the final property of the optimal code with the minimal conductance.

#### Lemma 5.

The best genetic code in terms of its maximum conductance must be determined by a partition of codon groups in which there are only groups of the size k = 3 and k = 4 with the minimal conductance, i.e. ϕ_3_(*G*) and ϕ_4_(*G*), respectively. Such code has only one codon group of the size k = 4.

*Proof.* It is an immediate conclusion from the observations presented above.

The example of the genetic code structure fulfilling Lemma 5 is presented in Fig. 4a. Its conductance is 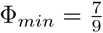. This structure consists of one fourfold degenerated group of codons and twenty groups of threefold degenerated codons.

**Figure 4.**
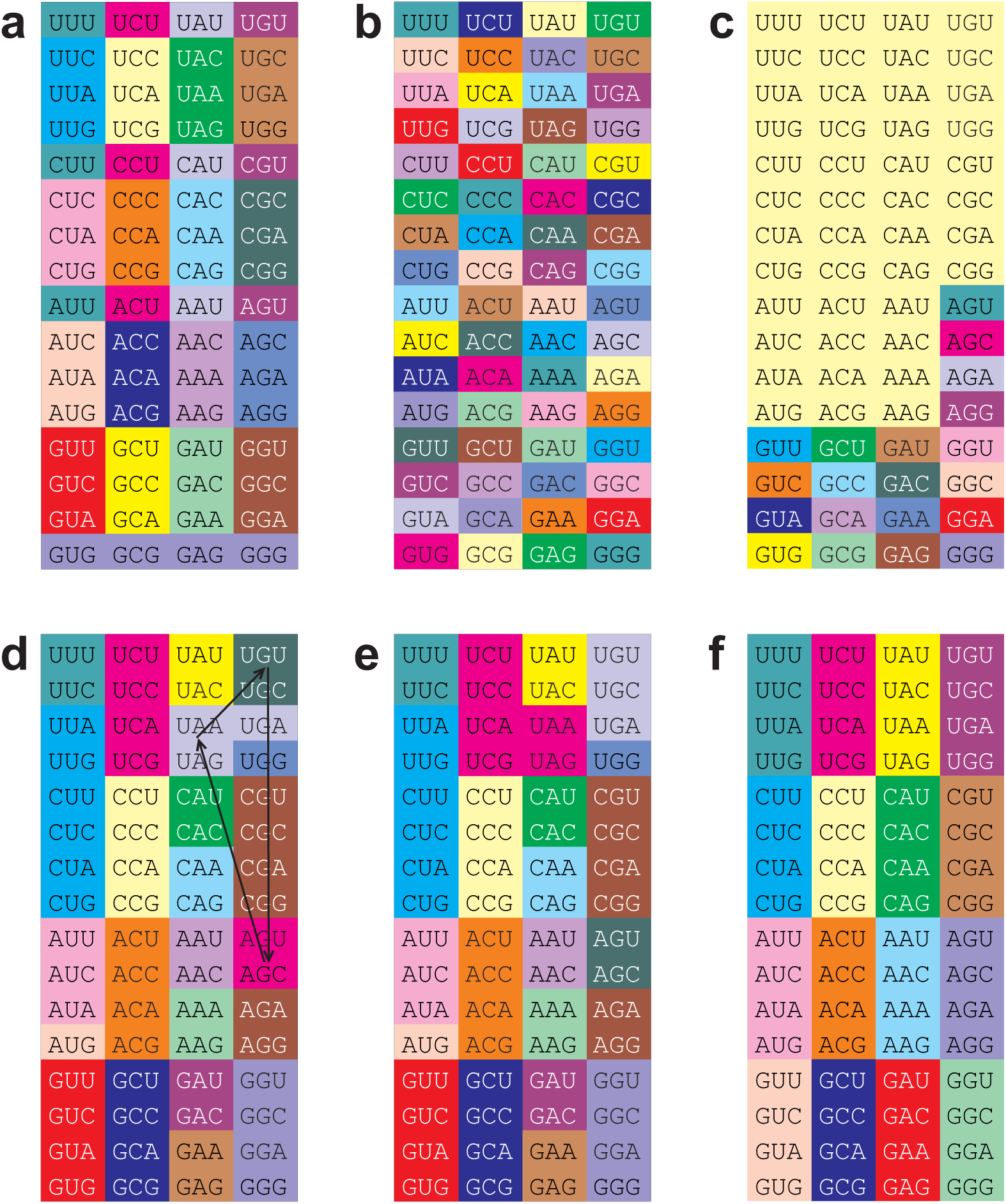
Various structures of genetic codes encoding 21 items and showing interesting properties in terms of code conductance. (a) An example of code that shows the minimum of the maximum code conductance Φ_*min*_ and simultaneously the minimum of the average code conductance 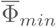. (b) An example of code showing the largest possible maximum and average conductance 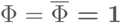 and consisting of one fourfold degenerated codon group and twenty groups of threefold degenerated codons, as the code presented in (a). (c) An example of code that shows the largest 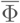 and consists of codon groups, each with its *k*-size-conductance *ϕ*_*k*_(*G*). (d) The standard genetic code (SGC). The arrows show the minimum number of changes in the SGC to obtain the best code in the SGC equivalence class with the *k*-size-conductance *ϕ*_*k*_(*G*). This code is presented in (e). (f) An example of code that encodes 16 items and shows the minumum of the maximum code conductance Φ_*min*_.

### 1.3 The optimal genetic code with respect to average conductance

An alternative approach to minimizing the maximum conductance of codon groups is based on minimizing the average conductance of a code. This measure admits a wider range of values of clusterings. We prove that the minimum value of average conductance achieved by a clustering of a codon graph into 21 groups is 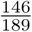. We begin with lemma that gives us a lower bound for the average code conductance calculated for any clustering of *G* in which the maximum size of codon groups is less or equal to 9.

#### Lemma 6.

Any clustering of G into 21 groups of sizes at most 9 has average conductance at least 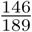.

*Proof.* Consider the following primal linear program computing a lower bound on the total conductance, (i.e. the average conductance multiplied by the number of groups, 21), of any code consisting of codon groups of sizes not bigger than 9. In the primal linear program, variables *x*_*i*_ correspond to the relaxed numbers of groups of size *i*, for 1 ≤ *i* ≤ 9; here “relaxed” means that these numbers are not necessarily integers, although their range is in [0, 21]. Note that since we are deriving a lower bound, if it holds for relaxed variables *x*_*i*_ denoting the number of groups of size *i* in the optimal solution, it automatically holds in the case when *x*_*i*_ are integers.

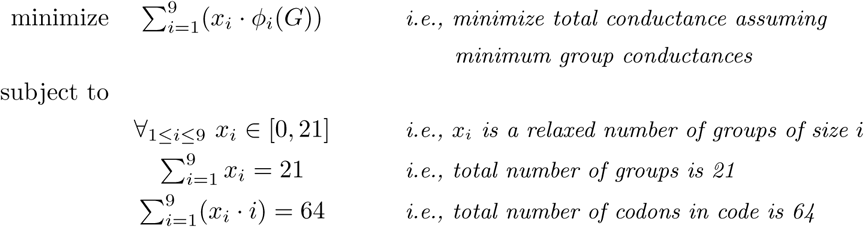

The dual of the primal program presented above is as follows:

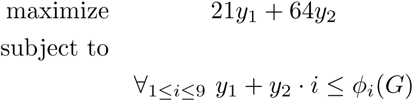

Proposition 4 guarantees that the values *ϕ*_*i*_(*G*), used in this linear program and taken from Fig, 2 are correct lower bounds of conductances of clusters of sizes up to 9,

It is easy to check, by a straightforward calculation, that the solution to the primal program is not greater than 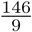, by setting *x*_3_ = 20, *x*_4_ = 1 and all other values of *x*_*i*_ to zeros. Similarly, the solution to the dual program is not smaller than 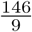, by putting 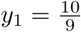 and 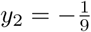. By the weak duality theorem [Cormen et al., 2009], the solution to the primal program is not smaller than the solution to the dual; hence we get that the solutions to these two programs must be equal and are exactly 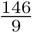. Therefore, for any possible combination of (integer) cluster numbers *x*_*i*_, the resulted total conductance is at least 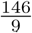. Thus, the minimum average conductance of a code is at least 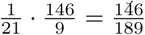.

Next we prove that the clustering of *G* into *k* = 21 groups minimizing the average conductance cannot contain a group of size bigger than 9.

#### Lemma 7.

No clustering of G into 21 clusters with a group size bigger than 9 has the average conductance smaller than 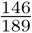.

*Proof.* The proof is by contradiction. Suppose, to the contrary, that there is a clustering of *G* into 21 groups that minimizes the average conductance and has group(s) of size bigger than 9. There are only four cases possible, described below and parametrized by 1 ≤ *j* ≤ 4:

**Case** *j***, for** 1 ≤ *j* ≤ 4: There are exactly *j* groups of size bigger than 9.

In the case *j*, the other 21 *− j* groups are selected out of at most 64 − 10 *j* codons.

Note that the cases for *j ≥* 5 are not feasible, as for *j* = 5 the number of codons in the groups of size smaller than 10 would be at most 64 10 5 = 14 and they should be distributed into 21 − 5 = 16 groups, which is impossible; the cases for *j ≥* 6 are infeasible by similar arguments.

For each of the four feasible cases, for 1 ≤ *j* ≤ 4, consider the following primal linear program computing a lower bound on the total conductance of any clustering of at most 64 − 10*j* codons into 21 *− j* groups of sizes not bigger than 9, in which variables *x*_*i*_ correspond to the relaxed numbers of clusters of size *i*, for 1 ≤ *i* ≤ 9; similarly as in the proof of Lemma 6 “relaxed” means that these numbers are not necessarily integers, although their range is in [0, 21 *− j*].

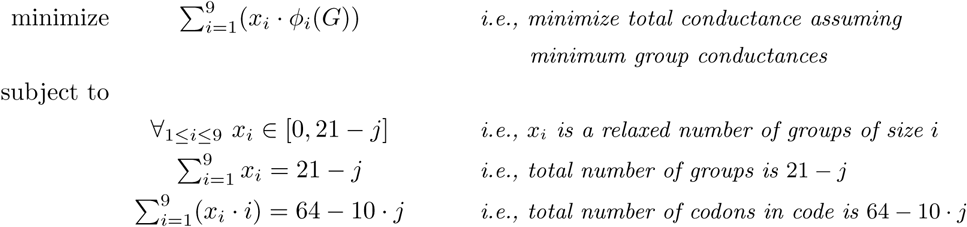

The dual of the primal program presented above is as follows:

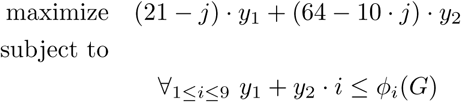

Proposition 4 guarantees that the values *ϕ*_*i*_(*G*), used in this linear program and taken from Fig, 2 are correct lower bounds on conductances of clusters of sizes up to 9.

It is easy to check, by a straightforward calculation, that the solution to the primal program is not greater than:

**for** 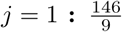, by setting *x*_3_ = 14, *x*_2_ = 6 and all other values of *x*_*i*_ to zeros;

**for** 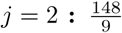, by setting *x*_3_ = 4, *x*_2_ = 15 and all other values of *x*_*i*_ to zeros;

**for** 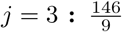, by setting *x*_2_ = 16, *x*_2_ = 2 and all other values of *x*_*i*_ to zeros;

**for** 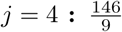, by setting *x*_3_ = 7, *x*_2_ = 10 and all other values of *x*_*i*_ to zeros.

Similarly, the solution to the dual program is not smaller than 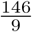, by putting 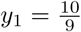 and 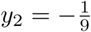, for every 1 ≤ *j* ≤ 4. By the weak duality theorem [Cormen et al., 2009], the solution to the primal program is not smaller than the solution to the dual; hence we get that the solutions to the primal program must be not smaller than 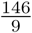. Therefore, for any possible combination of (integer) cluster numbers *x*_*i*_, the resulted total conductance is at least 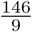 in all four cases. Hance, the average conductance of the whole clustering is bigger than 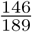 in all four cases. □

#### Theorem 1.

A clustering of G into 21 clusters that minimizes the average conductance achieves the value 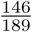.

*Proof.* From Lemma 6, any clustering into groups of size at most 9 has the average conductance of at least 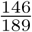. By Lemma 7, no clustering of *G* into 21 groups with a group of size bigger than 9 has the average conductance smaller than 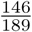. On the other hand, there is a clustering into twenty groups of size 3 and one group of size 4 such that each group of size 3 has conductance 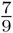 and the group of size 4 has conductance 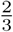, resulting in the average conductance of the clustering equal to 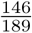. It can be checked that the clustering presented in Fig. 3 has the abovementioned properties. In view of the two cited lemmas, this clustering achieves the minimum possible value of the average conductance.

### 1.4 The general properties of genetic codes in respect to the average *κ*-size-conductance

In the previous section we gave a lower bound of the average code conductance but it would be interesting to determine some general properties of the optimal genetic codes in terms of this measure. To deal with this problem, we apply the Definition 6 of the average *κ*-size-conductance and the Proposition 1. As a consequence, we get another way to prove the Theorem 1 because it is enough to calculate the average *κ*-size-conductance for all possible equivalence classes [*κ*]. This calculation is possible by using the Proposition 4 because it gives us a way to compute the exact value of *ϕ*_*k*_(*G*) for each *k ≥* 1. Therefore, we are able to calculate the average *κ*-size-conductance for all [*κ*]. We evaluated the value of Φ[*κ*] for all 59, 755 equivalence classes defined by vectors of integers *κ* under the condition (1). All these cases are presented in Electronic Supplementary Material in ESM_2. Note that the number 59, 755 is equal to the number of monotonic representations of 64 by 21 non-zero summands, *P* (64, 21). Basing on these data, we can formulate the subsequent propositions:

#### Proposition 5.

The average κ-size-conductance of any code is not smaller than 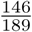.

This proposition corresponds to the thesis of Theorem 1. The next proposition gives us another feature of the optimal genetic code.

#### Proposition 6.

There are only 44 equivalence classes [κ] for which the average κ-size-conductance is equal to 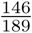. Moreover, for these [κ] classes, we found at least one partition 𝒞 of the graph G which fulfills the condition 𝒞 ∈ [κ]. As a consequence the equality 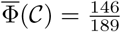 holds.

The last proposition states a very interesting characteristic of the optimal graph *G* partition in terms of the average code conductance and is an improvement of the theoretical result of Lemma 6. Note that we obtained it by using a computational support, which implements and analyzes the abovementioned (formally justified) argumentation, c.f., the Electronic Supplementary Material in ESM_2.

#### Proposition 7.

Let S_max_ = max_S∈𝒞_ #S be the maximum size of a codon group which belongs to the partition 𝒞. Then for every partition 𝒞 of the graph G into 21 sets, 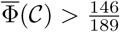, if S_max_ > 4.

In other words, there exists no optimal genetic code, in terms of minimizing the average code conductance, in which *S*_*max*_ > 4. This proposition follows directly from the Proposition 1 and the fact that the respective average *κ*-size-conductance for *κ* = (*k*_1_, *k*_2_, *…, k*_21_), where max_*i*_ *k*_*i*_ > 4, achieves greater values than 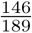, c.f., the Electronic Supplementary Material in ESM_2.

It is also interesting that the best code in terms of minimizing the average code conductance and the maximum conductance, presented in Fig. 4a, as well as the worst code maximizing these parameters, shown in Fig. 4b, belong to the same equivalence class.

#### The properties of the standard genetic code in terms of conductance

It is evident, that the standard genetic code (SGC) is far from being optimal in terms of the code maximum conductance Φ(*𝒞*) because this parameter for the standard genetic code equals 1, which is the worst possible value. This is the consequence of the fact that the standard genetic code contains two codon groups consisting of only one codon. The codon group {*AUG*} encodes methionine and the group {*UGG*} encodes tryptophan. Each single-nucleotide substitution in these codons causes the change in the translation of the protein-coding sequences.

The performance of the SGC changes when we investigate its average code conductance. The value of 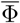(*SCG*) is equal to 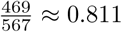, which is definitely closer to the optimal solution 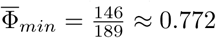 (Fig. 4a) than to the largest possible average conductance that equals 1 (Fig. 4b). Moreover, Φ(*SGC*) is also smaller than the average conductance 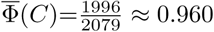 calculated for the worst code consisting of codon groups optimal in terms of *k*-size-conductance *ϕ*_*k*_(*G*).

Moreover, the SGC is quite good in its own equivalence class of codes because the average *κ*-size-conductance of the best code in this class is 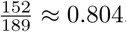, i.e. is only slightly lower than 0.811 (Fig. 4d and e). The SGC performs also well in the general comparison with all possible 59, 755 equivalence classes of codes. Assuming that for all these classes, it is possible to find at least one representative with its average *κ*-size-conductance, there are only 2778, i.e. 4.6% of cases with the 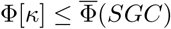. The average conductance of the SGC is located at the left tail of the Φ[*κ*] distribution (Fig. 5).

**Figure 5.**
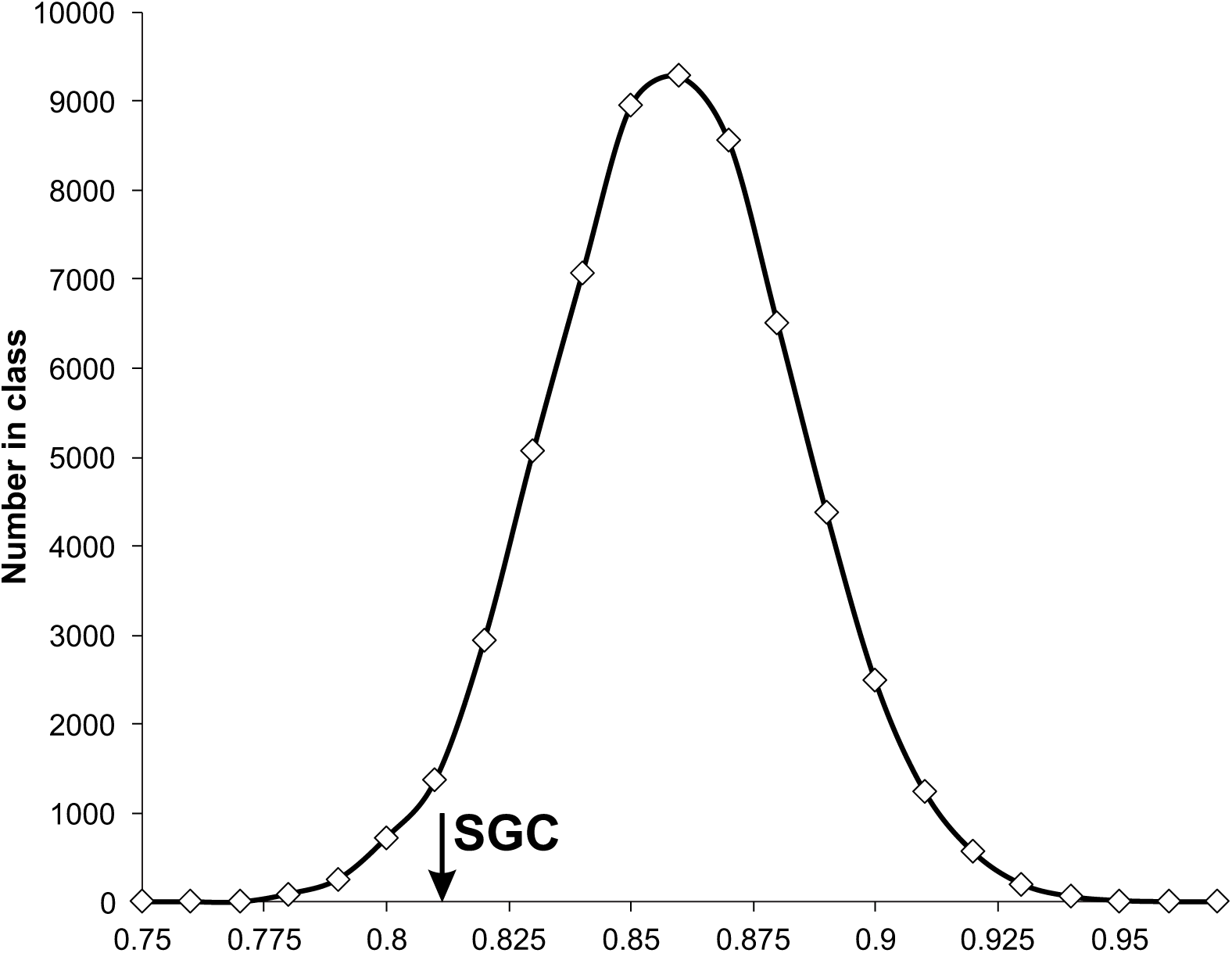
The distribution of Φ[*κ*] values calculated for all possible 59, 755 equivalence classes of codes. The value of the standard genetic code (SGC) is indicated by the arrow.

In fact, the SGC has many codon groups optimal in terms of the *k*-size-conductance (Table 2). All groups of fourfold degenerated codons have the minimal conductance *ϕ*_4_(*G*) for their size. Similarly, the codon groups of twofold degenerated codons also show the minimal conductance *ϕ*_2_(*G*) for their size. However, the conductance of the codon groups with the size *k* = 3 and *k* = 6 is more diversified. There are two groups consisting of three codons. One encodes isoleucine and the other stop translation signal. The isoleucine codon group has the minimal conductance 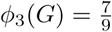 for its size, whereas the conductance of the stop codon group is not optimal:

**Table 2.** The structure of the standard genetic code in terms of the codon groups conductance. Each row describes: the amino acid encoded by the respective codon group, the size of the codon group, its conductance *ϕ*(*S*) and *ϕ*_*k*_(*G*), i.e. the minimal conductance of the codon group with the size *k*.

| AA | Codon group ($S$) | Size $k$ | $\phi(S)$ | $\phi_k(G)$ |
| --- | --- | --- | --- | --- |
| Ala | $\{GCA, GCU, GCG, GCC\}$ | 4 | $\frac{2}{3}$ | $\frac{2}{3}$ |
| Arg | $\{AGA, AGG, CGA, CGU, CGG, CGC\}$ | 6 | $\frac{2}{3}$ | $\frac{2}{3}$ |
| Asn | $\{AAU, AAC\}$ | 2 | $\frac{8}{9}$ | $\frac{8}{9}$ |
| Asp | $\{GAU, GAC\}$ | 2 | $\frac{8}{9}$ | $\frac{8}{9}$ |
| Cys | $\{UGU, UGC\}$ | 2 | $\frac{8}{9}$ | $\frac{8}{9}$ |
| Gln | $\{CAA, CAG\}$ | 2 | $\frac{8}{9}$ | $\frac{8}{9}$ |
| Glu | $\{GAA, GAG\}$ | 2 | $\frac{8}{9}$ | $\frac{8}{9}$ |
| Gly | $\{GGA, GGU, GGG, GGC\}$ | 4 | $\frac{2}{3}$ | $\frac{2}{3}$ |
| His | $\{CAU, CAC\}$ | 2 | $\frac{8}{9}$ | $\frac{8}{9}$ |
| Ile | $\{AUA, AUU, AUC\}$ | 3 | $\frac{7}{9}$ | $\frac{7}{9}$ |
| Leu | $\{UUA, UUG, CUA, CUU, CUG, CUC\}$ | 6 | $\frac{2}{3}$ | $\frac{2}{3}$ |
| Lys | $\{AAA, AAG\}$ | 2 | $\frac{8}{9}$ | $\frac{8}{9}$ |
| Met | $\{AUG\}$ | 1 | 1 | 1 |
| Phe | $\{UUU, UUC\}$ | 2 | $\frac{8}{9}$ | $\frac{8}{9}$ |
| Pro | $\{CCA, CCU, CCG, CCC\}$ | 4 | $\frac{2}{3}$ | $\frac{2}{3}$ |
| Ser | $\{AGU, AGC, UCA, UCU, UCG, UCC\}$ | 6 | $\frac{40}{54}$ | $\frac{2}{3}$ |
| Thr | $\{ACA, ACU, ACG, ACC\}$ | 4 | $\frac{2}{3}$ | $\frac{2}{3}$ |
| Trp | $\{UGG\}$ | 1 | 1 | 1 |
| Tyr | $\{UAU, UAC\}$ | 2 | $\frac{8}{9}$ | $\frac{8}{9}$ |
| Val | $\{GUA, GUU, GUG, GUC\}$ | 4 | $\frac{2}{3}$ | $\frac{2}{3}$ |
| Stp | $\{UAA, UAG, UGA\}$ | 3 | $\frac{23}{27}$ | $\frac{7}{9}$ |

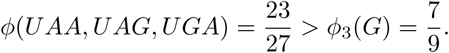

Considering the codon groups with the size *k* = 6, those encoding arginine and leucine have the minimal conductance 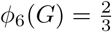 for their size, whereas the codon group for serine is not optimal in terms of the conductance minimization because it can be described by the following inequality:

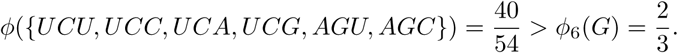

To summarize, the properties of the standard genetic code in terms of the conductance measure lead to ambiguous conclusions. On the one hand, this code is the worst according to its Φ(*𝒞*). It is also not optimal in terms of the average conductance. Moreover, in both cases it could be improved just by small number of changes. On the other hand, out of 19 codon groups with more than one codon, 17 show the *k*-size-conductance for their size.

If we assume that the standard genetic code evolved to minimize the costs of mutations and translation errors [Ardell, 1998, Di Giulio, 1989, Freeland and Hurst, 1998b, Freeland and Hurst, 1998a, Freeland et al., 2003, Haig and Hurst, 1991, Woese, 1965], then the lack of its full optimization, in terms of the code conductance and the average code conductance, can result from its stepwise evolution. It seems probable that the present form of the standard genetic code evolved from a code encoding a smaller number of amino acids [Di Giulio, 2008, Higgs and Pudritz, 2009, Massey, 2016, Sun and Caetano-Anollés, 2008]. Therefore, if the process of optimization occurred at subsequent stages of code evolution then the structure that appeared at a given stage did not have to be optimal in the next stage after the addition of other amino acids. What is more, after the expansion of the code, the full re-optimization might not have been possible because it would have caused changes in the translation of codons to amino acids and consequently, dramatic changes in many sequences of already encoded proteins. Such evolving code inherited the fixed assignments of codons to amino acids from previous stages and the final form of the code does not have to be optimal in general. For example, let us consider a simple optimal code with the code conductance 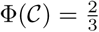 encoding fifteen amino acids and stop translation signal by sixteen codon groups with the minimal conductance (Fig. 4f). To obtain the optimal code which encodes 21 amino acids and stop signal with the code conductance 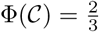, it is sufficient to add only five amino acids but it would result in substantial changes in as many as 15 codon groups. It is evident that the evolution from the optimal code at a given stage to the optimal code at the next stage would require many fundamental changes not only in the assignments of codon groups but also in the translated polypeptides.

Since the standard genetic code does not seem to be fully optimized to minimize the effects of mutations or translational errors because much better codes can be found [Blazżej et al., 2016, Novozhilov et al., 2007, Santos et al., 2011, Santos and Monteagudo, 2017], other factors must have taken part in shaping its structure as well. The addition of subsequent amino acids into the standard code could have proceeded according to their relationships in biosynthetic pathways as claims the co-evolution theory [Di Giulio, 1997, Di Giulio and Medugno, 1999, Di Giulio, 2004, Di Giulio, 2008, Wong, 1975, Wong et al., 2016]. Consequently, the potential tendencies of this code to minimize the errors may be a by-product of this process [Di Giulio, 2016, Di Giulio, 2017]. Other studies have also showed that no direct selection for the error minimization was necessary to produce the genetic codes with this property, which could have evolved as a result of gene duplications of adaptors and charging enzymes [Massey, 2015, Massey, 2016]. Interestingly, the optimization of biological systems to minimize the harmful effects of mutations does not have to require changes in the genetic code because the mutational pressure can be subjected to this optimization around the fixed genetic code [Dudkiewicz et al., 2005, Mackiewicz et al., 2008, B laz˙ej et al., 2015, B laz˙ej et al., 2017].

## Conclusions

Our results show that the general structure of genetic code and the problem of the genetic code optimality can be successfully reformulated using a methodology adapted from graph theory in the context of optimal clustering of a specific graph. To evaluate the quality of the genetic code, we defined the code maximum conductance and the average code conductance. The former evaluates a given genetic code in terms of its “weakest link”, i.e. the codon group with the maximum set conductance, whereas the latter takes into account the values of all codon groups of the code. From the biological point of view, these two measures describe the code robustness against amino acid and stop signal replacements resulting from single nucleotide substitutions between codons. According to this relatively general assumptions, we found the optimal code that minimizes its code conductance and differs from the standard genetic code although the SGC has many optimal codon groups with the minimal conductance for their size. It implies that the role in the organization of the genetic code was played not only by the selection for the minimization of amino acid and stop signal replacements but also by the stepwise evolution of the code associated with its expansion and addition of subsequent amino acids, e.g. according to the evolution of biosynthetic pathways.

## Electronic Supplementary Material

1. ESM_1. The adjacency matrix of the codon graph *G*, where rows and columns are ordered in lexicographical order. Zero values were omitted for clarity.
2. ESM_2. All possible 59, 755 equivalence classes of codes coding 21 items by 64 codons, with the average *κ*-size-conductance.

